# Neural correlates of phenomenological attitude toward perceptual experience

**DOI:** 10.1101/2024.07.07.602347

**Authors:** Satoshi Nishida, Hiro Taiyo Hamada, Takuya Niikawa, Katsunori Miyahara

**Affiliations:** Center for Information and Neural Networks (CiNet), Advanced ICT Research Institute, National Institute of Information and Communications Technology (NICT), 1-4 Yamadaoka, Suita, Osaka 565-0871, Japan; Graduate School of Frontier Biosciences, Osaka University, 1-3 Yamadaoka, Suita, Osaka 565-0871, Japan; Neurotechnology R&D Unit, Araya Inc., 1-12-32 Akasaka, Minato-ku, Tokyo 107-6024, Japan; Department of Functional Brain Imaging, National Institute of Radiological Sciences, National Institutes for Quantum Science and Technology, 4-9-1 Anagawa, Inage-ku, Chiba 263-0024, Japan; Neural Computation Unit, Okinawa Institute of Science and Technology, Graduate University, 1919-1 Tancha, Onna-son, Okinawa 904-0495, Japan; Graduate School of Humanities, Kobe University, 1-1 Rokkodai-machi, Nada-ku Kobe, Hyogo 657-8501, Japan; Center for Human Nature, Artificial Intelligence, and Neuroscience (CHAIN), Hokkaido University, Kita 12 Nishi 7, Kita-ku, Sapporo, Hokkaido, 060-0812, Japan

**Keywords:** Phenomenological reduction, Conscious experience, Human brain, fMRI

## Abstract

Phenomenology is one of the most promising approaches to study conscious experience. It holds that a rigorous study of conscious experience requires a transition in the subject from the “natural attitude” (NA) to the “phenomenological attitude” (PA). NA describes our ordinary stance, in which our attention is directed at external objects and events. PA is a distinctive, reflective stance in which our attention is directed at our conscious experience itself. Despite its theoretical importance in philosophy and science of consciousness, the neural mechanisms underlying PA remain unknown. To clarify this point, we developed a novel behavioral task in which participants alternate between NA and PA in relation to their stimulus-evoked subjective experiences. Participants are presented with two sentences and requested to identify the one that best captures their experience. These sentences are designed to induce either NA or PA. We found that participants had lower error rates but slower reaction times in the PA condition compared to the NA condition, suggesting a difference beyond task difficulty. Using fMRI, we also found that multivoxel activation patterns in the premotor cortex, posterior parietal cortex, supplementary motor area, and cerebellum successfully classified the task conditions. Furthermore, the activation strength in these regions was lower in the PA condition, indicating that PA depends on neural processes that suppress action-related information. These findings provide the first evidence for the neural signature of PA, contributing to a better understanding of phenomenological method and its underlying neural mechanisms.

**Significance statement:** Phenomenology is one of the most promising approaches to study conscious experience. A key step is a transition from the natural attitude (NA)—where our attention is directed at external objects and events—to the phenomenological attitude (PA)—where our attention is directed toward our conscious experience itself. However, the neural processes underlying PA remain unclear. This study aimed to clarify this point by analyzing fMRI signals measured during a cognitive task that forced participants to repeatedly alternate between NA and PA. We observed that multiple action-related brain regions exhibited different activation patterns between NA and PA. Our findings provide the first neuroscientific evidence that illuminates the core process of the phenomenological method.

## Introduction

Phenomenology is considered one of the most promising approaches for studying conscious experience. Originally developed by Edmund Husserl in the late 19th century as an innovative philosophical method, it has become one of the most influential philosophical approaches in the 20th century and into the present day. Phenomenology emphasizes the value of investigating structures of consciousness, such as intentionality, temporality, and minimal self-awareness, from a first-person perspective (1–3). Thus, phenomenologists study consciousness first and foremost by reflecting upon their concrete experience, rather than constructing hypotheses based on commonsense intuitions and metaphysical speculations.

Phenomenologists hold that a rigorous study of conscious experience requires a methodological operation called the “phenomenological reduction,” which involves transition from the “natural attitude” (NA) to the “phenomenological attitude” (PA). NA describes our ordinary stance, in which our attention is directed at external objects and events, such as the moth resting on the window, the meeting starting soon, etc. To investigate the structures of consciousness, however, we need to focus less on these objects of consciousness (or what we are conscious of) than on the conscious experience as such (or how we are conscious of these objects). PA refers to this alternative, reflective stance whereby our attention is directed at our own conscious experience.

Recently, the phenomenological method is gaining significant traction thanks to the resurgence of the science of consciousness. The ultimate goal of consciousness science lies in understanding the physical mechanism underlying conscious experience; many researchers have recognized that this requires understanding the structures of the conscious experience itself (4). In cognitive science, however, there is no established method to specifically uncover the structures of consciousness. Phenomenology could be a promising approach to fill this gap in research (5–7).

Nonetheless, the nature of PA is hardly explored in the cognitive sciences, despite its importance both in classical phenomenology and contemporary consciousness research. What are the underlying neural mechanisms of PA? What cognitive and psychological functions does the transition from NA to PA include? What can we learn about the nature of these attitudes from their behavioral and neural signatures? These questions have not been investigated so far.

To address these questions, we developed an original behavioral task, the natural– phenomenological attitude switching (NPAS) task, where participants are led to alternate between NA and PA in relation to stimulus-evoked subjective experiences from trial to trial (Fig. 1). In each task trial, participants are asked to determine which of the two sentences correctly describes a visual-scene stimulus. The two sentences are designed to automatically induce either NA or PA in the participant under NA and PA conditions, respectively. In NA conditions, participants are guided to attend to the objects depicted in the image. In contrast, in the PA conditions, participants are guided to attend to the horizonal structure of their visual experience.

**Figure 1.**
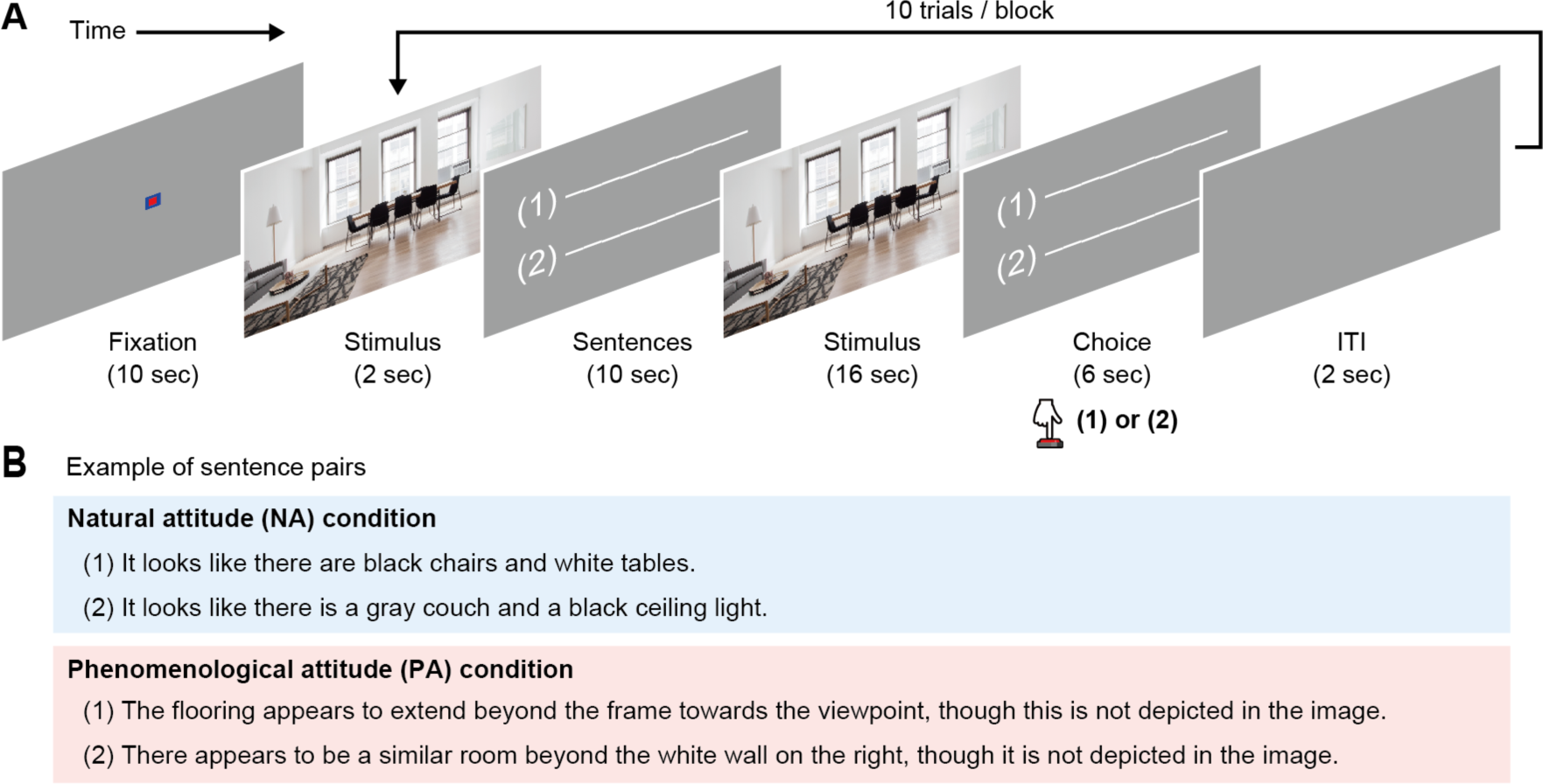
Natural–phenomenological attitude switching (NPAS) task. (A) Task procedure. Each block started with a 10-s fixation period, followed by 10 trials of the NPAS task. In each trial, after a 2-s presentation of an image stimulus, a pair of sentences was displayed for 10 s. Either sentence correctly described the content of the participant’s perceptual experience evoked by the image stimulus. Then, after a 16-s reappearance of the stimulus, the sentence pair was displayed again for 6 s (choice period). Participants had to determine which sentence was correct and respond within this choice period using a button press. (B) Example of sentence pairs. An NA pair, which made participants perceive the image in NA (top), and a PA pair, which made them perceive the image in PA (bottom), are separately shown. These pairs were both associated with the image stimulus in A and presented with the same stimulus but in a separate block of the NPAS task. Sentence (1) in each pair is the correct answer for this stimulus.

The “horizon” is a technical term in phenomenological philosophy that refers to aspects of conscious experience that exceed the objects of experience of which the subject is thematically aware [pp. 162–163 in (2)]. For example, when I see an apple on a table, I am most vividly aware of the side of the apple facing to me. Yet at the same time, I am also indirectly aware of the backside of the apple or the part of the table under the apple from which I receive no visual input. I am aware that they are there and that they can be presented in further acts of perception if I pick up the apple and turn it around. Without this horizonal awareness, it would be impossible to experience “an apple on a table,” a state that constitutively implies the existence of the backside of the apple or the hidden part of the table supporting it. The fact that conscious experience always involves such a horizonal structure is widely regarded as a major discovery of phenomenological philosophy––that is, something explicit only after Husserl and other phenomenologists approached conscious experience from PA.

We examined the behavioral and neural signature of PA by analyzing differences in behavior and functional magnetic resonance imaging (fMRI) signals depending on task conditions.

## Results

### Behavioral performance

To gain insights into the behavioral signature of PA, we measured the error rate and reaction time of participants in the NPAS task. Figure 2 displays the error rate and reaction time separately for the NA and PA conditions of the task. We found a lower error rate in the PA condition (NA: mean = 0.16, 95% confidence interval [CI] = 0.14–0.19; PA: 0.92, 0.79–0.11; Wilcoxon signed-rank test, P < 10^−7^). Yet, the reaction time was longer in the PA condition (NA: mean = 2.43 s, 95% CI = 2.32– 2.56 s; PA: 2.77 s, 2.62–2.93 s; P < 10^−6^). Additionally, we observed a strong correlation between error rate and reaction time in PA trials (Pearson r = 0.75, P < 10^−7^) but no significant correlation in NA trials (r = 0.17, P = 0.29; Fig. 2C). These results suggest that participants employed different behavioral strategies for NA and PA conditions, rather than using the same strategy for two kinds of tasks with varying levels of difficulty.

**Figure 2.**
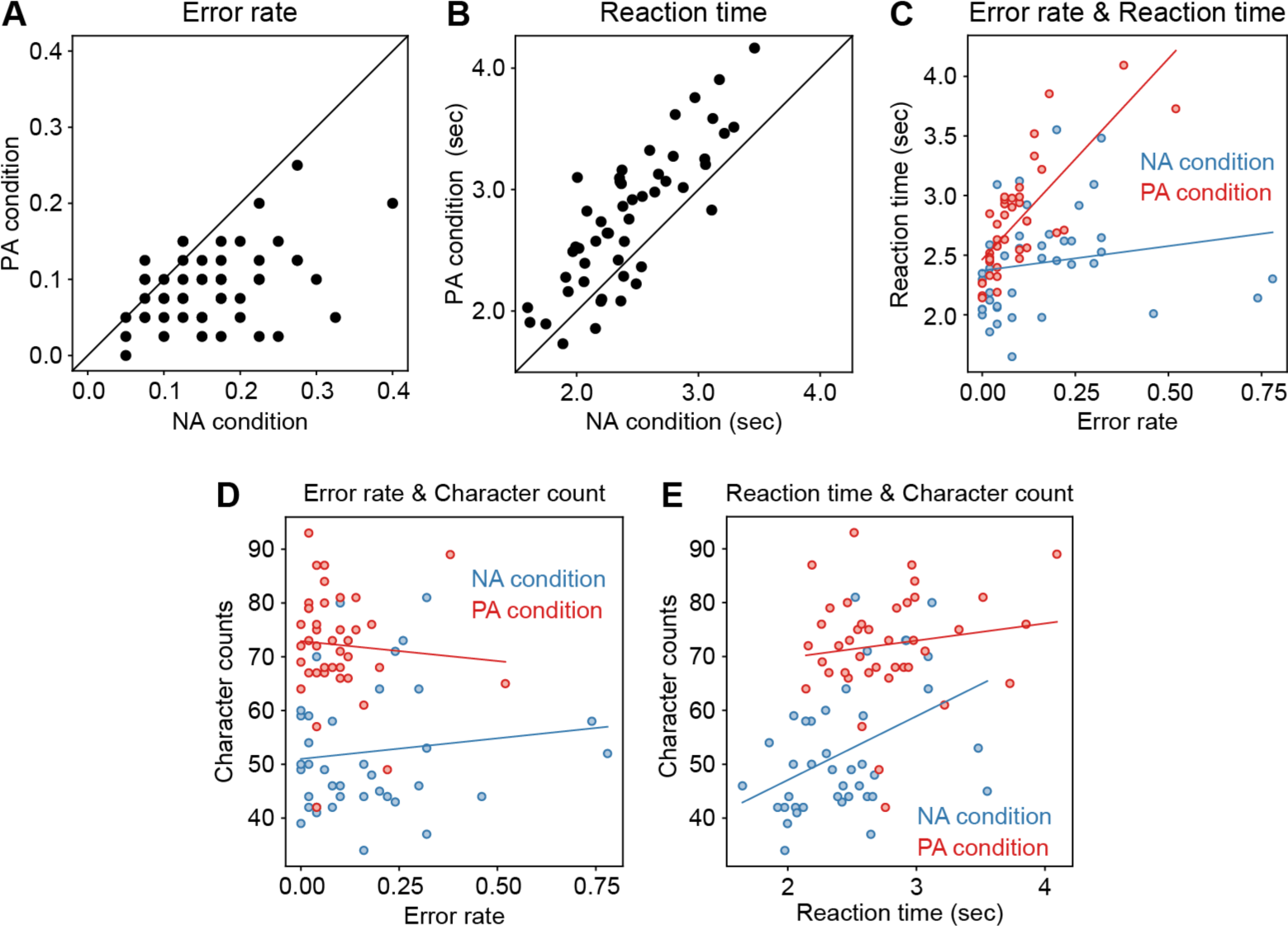
Differences in behavioral characteristics between NA and PA conditions in the NPAS task. (A–B) Behavioral comparison between conditions. The error rate (A) and mean reaction time (B) for each participant, represented by each dot, were compared between NA (x-axis) and PA (y-axis) conditions. (C) Correlation between error rate (x-axis) and reaction time (y-axis) in each condition (blue, NA; red, PA). These measures were evaluated for each stimulus-sentence pair from all participants. Each dot represents a single pair. Blue and red lines denote linear fits to the data for NA and PA conditions, respectively. (D–E) Association between behavioral measures (D, error rate; E, reaction time) and character counts in individual sentence pairs for each condition (blue, NA; red, PA). Character counts were added within a pair of sentences for each stimulus. Other conventions are the same as in (C).

The character count for each pair of sentences was higher in PA sentences (mean ± SD = 35.2 ± 5.60 Japanese characters) compared to NA sentences (mean ± SD = 25.9 ± 5.95 Japanese characters; t-test, P < 10^−18^). However, the observed differences in error rate and reaction time (Fig. 2) were unlikely to be due to character count differences. The average error rate for participants in each pair of stimuli and sentences did not significantly correlate with the character counts of those pairs in both NA and PA conditions (Fig. 2D; NA: Pearson r = 0.12, P = 0.47; PA: r = −0.075, P = 0.64). Additionally, the average reaction time for participants significantly correlated with the character count in NA sentences but not in PA sentences (Fig. 2E; NA: r = 0.43, P < 0.01; PA: r = 0.14, P = 0.37). These results indicate that character count impacts reaction time only in the NA condition and does not affect other behavioral measures, suggesting little impact of character-count differences on condition-dependent changes in the behavioral measures.

### Whole-brain searchlight decoding

To investigate the brain regions that exhibit differential responses to stimuli based on NA and PA, we conducted whole-brain searchlight decoding. This technique entails the differentiation of NA and PA conditions based on multivoxel patterns of stimulus-induced fMRI signals within localized brain regions (for more details, see Materials and Methods). Our results revealed a significant decoding accuracy within a widely distributed network of regions (Fig. 3; P < 0.05, voxel-level family-wise error [FWE] corrected), including the premotor cortex (PMC), the posterior parietal cortex (PPC), the supplementary motor area (SMA), and the cerebellum. These observations suggest that these regions are implicated in distinct neural signaling between NA and PA.

**Figure 3.**
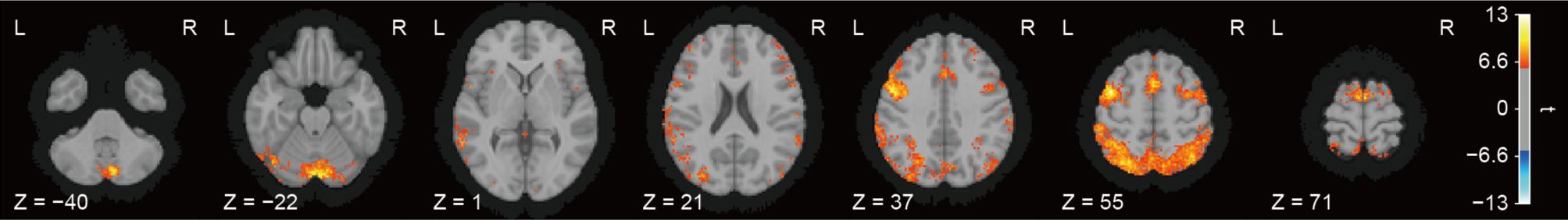
Whole-brain searchlight decoding to identify brain regions exhibiting different multivoxel activation patterns between NA and PA conditions. Colored locations indicate the centers of searchlights showing significant decoding accuracy (P < 0.05, voxel-level FWE corrected), denoted by t statistics, reflecting how accurately multivoxel patterns at the locations discriminate task conditions. No location showed significantly negative t statistics. L, left hemisphere; R, right hemisphere.

### Condition-dependent changes in activity strength

The decoding analysis provided evidence regarding the brain regions changing their activation patterns between NA and PA conditions. However, how the strength of activation changes depending on the conditions remains unclear. Therefore, we examined the variation in activation strength depending on the condition to determine whether PA enhances or suppresses activation in those brain regions.

Prior to this examination, we identified specific regions of interest (ROIs) for analysis from clusters of voxels showing significant decoding accuracy (see Materials and Methods). We identified a total of six ROIs, including the bilateral PMC, bilateral PPC, SMA, and cerebellum (SI Appendix, Fig. S1 and Table S1); these ROIs were used for subsequent analysis.

Figure 4 depicts the estimated activation strength in each ROI separately for the NA and PA conditions. In all ROIs, the activation strength was significantly lower in the PA condition than in the NA condition (Wilcoxon signed-rank test, P <10^−8^, ROI-level FWE corrected). This suggests that PA modulates activation in these regions in a suppressive fashion.

**Figure 4.**
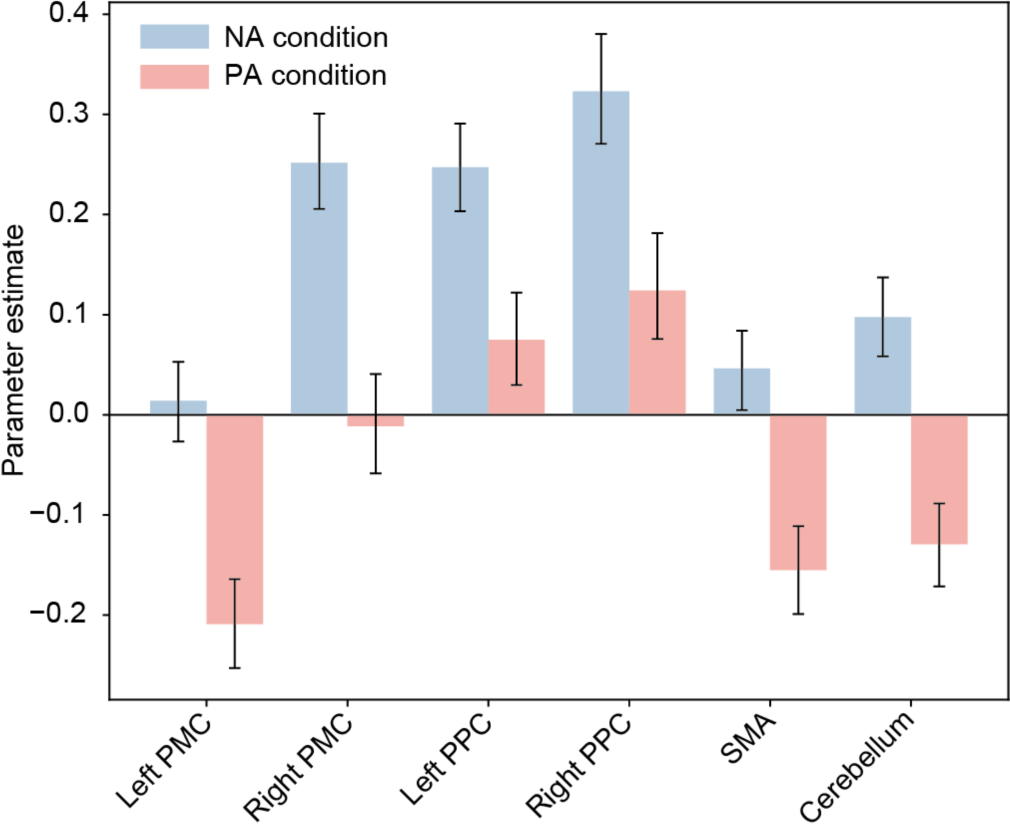
Condition differences in mean activation (parameter estimates) of the six ROIs. The six ROIs correspond to voxel clusters showing significant accuracies in searchlight decoding. The parameter estimates (z-scored β values estimated by a general linear model) among voxels within each ROI of each participant were averaged separately for each NA and (blue) and PA (red) condition. Then, ROI-wise parameter estimates were averaged across participants. Error bars denote 95% CI. The ROI-wise parameter estimates significantly differed between conditions in all ROIs (P < 10^-8^, Bonferroni corrected). PMC, premotor cortex; PPC, posterior parietal cortex; SMA, supplementary motor area.

### Potential confounding factors

The regions in which we observed successful decoding of task conditions (PMC, PPC, SMA, and cerebellum), are believed to play a role in action planning, production, and monitoring (8–11). Therefore, our observations may indicate activation-pattern changes within these regions depending on task performance, which varied between NA and PA conditions (Fig. 2A–B), rather than NA and PA alternation.

To test this possibility, we initially identified brain regions in which the reaction time could be decoded from brain activity. To achieve this, we employed another whole-brain searchlight decoding of fMRI signals to differentiate between trials with slower and faster reaction times (see Materials and Methods for more details). We found that only a small number of voxels exhibited significant decoding accuracy (SI Appendix, Fig. S2A; P < 0.05, voxel-level FWE corrected), unlike the decoding of task conditions (Fig. 3). Therefore, it is improbable that the observed decodability of PA and NA is due to trial-to-trial variation in reaction times.

We next explored the relationship between condition-dependent changes in brain activity and changes in error rates. If the observed decodability of task conditions was linked to changing error rates rather than the alternation of PA and NA, one would expect a correlation between decoding accuracy and condition-dependent changes in error rates. Specifically, a greater (or lesser) shift in error rates between the NA and PA conditions would be associated with higher (or lower) decoding accuracy. To test this, we calculated the difference in error rates between the PA and NA conditions for each participant and tested its correlation with decoding accuracy, averaged across participants for each of the six ROIs (SI Appendix, Fig. S1). However, our analysis revealed no significant correlation in any ROIs (SI Appendix, Fig. S3; Pearson r = −0.20–−0.046, P > 0.17, uncorrected). Thus, the changes in error rates dependent on conditions are unlikely to impact the decodability of PA and NA.

Finally, we explored the potential impact of character-count differences between PA and NA sentences. Even though these sentences were not shown during the stimulus presentation period used for decoding, it is possible that condition-dependent variations in character count might have influenced the decodability of PA and NA conditions. To test this, we conducted whole-brain searchlight decoding in fMRI signals for another time to differentiate between trials with more character counts and those with fewer character counts (see Materials and Methods for more details). We found that only a limited number of voxels exhibited significant decoding accuracy (SI Appendix, Fig. S2B; P < 0.05, voxel-level FWE corrected), in contrast to the decoding of task conditions (Fig. 3). This indicates that the character counts of the sentences had little impact on the observed decodability of PA and NA.

## Discussion

We investigated the behavioral and neural signatures of PA by analyzing changes in behavioral performance and brain activation patterns/magnitude between conditions in the NPAS task. We found that participants exhibited lower error rates and slower reaction times in the PA condition than in the NA condition. This difference indicates that these two conditions do not simply differ in terms of task difficulty. Task conditions were successfully classified using the multivoxel activation patterns in PMC, PPC, SMA, and the cerebellum, suggesting that these regions indicate differences between PA and NA. The activation strength in these regions was lower in PA conditions relative to NA conditions. To our knowledge, these findings provide the first evidence for the neural correlates of PA toward subjective experiences.

These results allow responding to skepticism on phenomenology (12), since critics doubt its effectiveness in achieving an epistemic effect. They might argue that PA is nothing but a theoretical artifact with no corresponding psychological reality. However, the current study indicates that the PA is a real psychological kind with an objectively measurable neural basis. Furthermore, it supports to the epistemic effectiveness of PA by showing that people with no previous exposure to phenomenological philosophy can discern the horizonal structure of conscious experience when appropriately guided into PA. These findings suggest that the phenomenological method is effective in uncovering the underlying structures of conscious experience.

Our behavioral data revealed different characteristics for NA and PA (Fig. 2). We speculate that this difference is attributed to additional cognitive processes required only for solving the task in the PA condition. In the NA condition, participants could read and understand the two sentences without difficulty choosing one of them based on their ability to match semantic contents with perceptual contents. This leads to a direct correlation between character count and reaction time simply because the longer the sentences, the longer it takes to read and comprehend them (Fig. 2E). Conversely, in the PA condition, participants needed to exercise additional cognitive processing, corresponding to the transition from NA to PA, to understand the sentences, which potentially required longer reaction times (Fig. 2B). However, once they understood the sentences, it was less challenging to solve the two-choice question in the PA condition than in the NA condition. Since comprehension difficulty does not necessarily match difficulty in selection, this could explain why participants showed a lower error rate in PA conditions (Fig. 2A).

To characterize the additional cognitive processes involved, we cross-referenced our neuroimaging data with existing models of PA. In the phenomenological literature, there are two prominent conceptual models of the transition from NA to PA, the Cartesian and the psychological way to phenomenological reduction, respectively (13, 14). The first highlights that NA involves a naïve belief in the existence of the world. It thus considers the transition from NA to PA in terms of the cognitive process of suspending one’s commitment to this naïve belief. Conversely, the second highlights the fact that NA involves practical interests in objects in the world. It thus envisions the transition to PA as involving a motivational process, whereby the practical interest in the world is inhibited. We found that participants adopting PA exhibit inhibited activity in brain regions associated with action production (i.e., PMC, PPC, SMA, and cerebellum). On the one hand, according to the Cartesian model, we should expect PA to be associated with the activity of brain regions related to higher-order cognition. On the other hand, if we follow the psychological model, which characterizes PA in terms of inhibition of practical interests, we should see inhibited activity in action-related regions of the brain. Accordingly, our neuroimaging data is more consistent with the psychological model. In this view, the additional cognitive process involved in PA conditions serves to inhibit their practical interests in the object characteristic of NA.

We could further clarify the nature of this cognitive process based on the concept of affordance. Affordance is a concept originally proposed by the ecological psychologist James J. Gibson (15). In its original meaning, it refers to the opportunities for actions available for an animal in a given environmental setting. Gibson claims that perception is primarily of affordances rather than objects and their intrinsic properties. Accordingly, we can interpret NA as an attitude invested with practical interests in affordances in the environment, which is inhibited in PA. Consistent with this interpretation, PA induced inhibited activity (PMC, SMA) and decreased activity (bilateral PPC) in brain regions related to the perception of affordances.^1^

The transition between NA and PA may involve switching between attentional sets (19). In the NA condition, participants focused their attention on object properties, while in the PA condition, they directed their attention toward the horizonal structure of conscious experience. Consequently, the NPAS task required alternating attentional sets between these two conditions. Different attentional sets are thought to change neural activation in the frontoparietal cortical network (19–21). In fact, a meta-analysis identifying brain regions involved in attentional switching underscored the importance of the premotor and posterior parietal cortices (22). Similarly, these regions exhibited differential activation patterns between NA and PA in our decoding analysis (Figs. 3 and SI Appendix, Fig. S1). Therefore, our findings could reflect changes in neural activation due to a previously unreported type of attentional difference, namely, between attention to external objects and one to conscious experience itself.

Could PA be characterized simply in terms of the exercise of visual mental imagery, rather than inhibition of practical interests? In the NPAS task, PA sentences typically referred to visual scenes not depicted in the stimulus image (e.g., a room beyond a wall, not shown within the image frame; see Fig. 1B and SI Dataset S1). In contrast, NA sentences generally referred to visual objects present in the stimulus images. Therefore, the different activation patterns between NA and PA could reflect activation changes corresponding to the involvement of visual mental imagery. However, in the phenomenological literature it is widely accepted that attending to perceptual experience and entertaining a visual mental imagery are different types of conscious acts: both involve sensory content, but the former is direct while the latter is indirect [pp. 120 in (1)]. Moreover, the brain regions exhibiting these differential activation patterns (Fig. 3 and SI Appendix, Fig. S1) did not include scene-selective cortical areas, such as the parahippocampal place area and the retrosplenial cortex. This result appears inconsistent with prior findings demonstrating an overlap in brain regions for both perception and mental imagery of the same information (23–25). Thus, it seems unlikely that visual mental imagery alone can explain our results.

Importantly, we found no brain region showing an enhanced activity during PA compared to its activity during NA (Fig. 4). This is initially puzzling, because some cognitive process appears to be necessary to initiate an attitude shift from NA to PA. In that case, we might expect to observe activities in the brain region that correlate with this cognitive operation in the neuroimaging data. We think the reason why we did not see this in our data is the task design rather than the nature of PA itself. We presented participants with the stimuli and the two alternative sentences before making a choice, while the neuroimaging data was obtained from the 6-s window when they made their choice (see Materials and Methods). In trials requiring PA, they would have likely began the attitude shift before their brain activity was recorded. Thus, the cognitive operation that initiates the attitude shift is not reflected in our neuroimaging data, which accounts for the absence of enhanced brain activity corresponding to PA. Correspondingly, the inhibition and decrease in brain activities differentiating PA from NA may reflect the subject’s state in PA, rather than the attitude shift that results in the adoption of PA.

One important limitation of our research is that it focused on the adoption of PA in relation to the horizonal structure of visual, perceptual experience. In the future, we will explore the neural mechanisms underlying PA in relation to other kinds of conscious experiences, such as other perceptual modalities (hearing, smelling, etc.), recollection, imagination, symbolic thinking, and action performance, since PA is considered domain-general, as it enables the subject to focus on their conscious experience regardless of the kind of conscious experience. The current experimental design allows to empirically approach this philosophical question by determining the brain activities corresponding to PA with respect to various kinds of conscious experiences. These future experiments could serve to validate the present results and support the domain-general conception of PA.

## Materials and Methods

### Participants

Fifty (22 females; age [mean ± SD] = 23.2 ± 4.1 SD years) healthy Japanese participants were recruited for the fMRI experiments. All had normal or corrected-to-normal vision. None had any experience studying phenomenology. Written informed consent was obtained from all participants and the experimental protocol was approved by the ethics and safety committees of NICT.

### Stimuli

Experimental stimuli consisted of 40 visual images collected from Pixabay (https://pixabay.com), a free stock photography and royalty-free stock media website. Twenty were natural images, and the remaining 20 were pictorial images (SI Dataset S1). The stimuli were presented full-screen at 4K (3840 × 2160) resolution (16.8 × 26.8 degrees of visual angle) on an MRI-compatible liquid crystal display (InroomViewingDevice, NordicNeuroLab, Norway).

### Task

In the fMRI experiments, each participant performed a natural–phenomenological attitude switching (NPAS) task where they had to alternate their NA and PA toward stimulus-evoked subjective experiences from trial to trial (Fig. 1). This task consisted of eight blocks, each starting with a 10-s fixation period, followed by 10 trials. This 10-s period served to avoid the undesirable effects of initial hemodynamic signal instability. In each trial, after a 2-s stimulus presentation, two sentences describing the stimulus were displayed vertically side by side for 10 s. Both sentences correctly explained the content of the participant’s subjective experience evoked by the stimulus. Then, after a 16-s stimulus reappearance, the two sentences were displayed again in the same order for 6 s (choice period). The participants were then asked to determine which sentence was correct and respond within this period by pressing either of two buttons with the index or middle finger of their right hand. The position (upper/lower) of the correct sentence was counterbalanced across trials.

Two kinds of sentence pairs were presented in each trial, NA pairs and PA pairs. One NA sentence and one PA sentence were assigned to each stimulus (Fig. 1B and SI Dataset S1); hence, each stimulus appeared twice in two separate trials with different pairs of sentences. The stimuli were presented in a random order, and NA and PA sentences were randomly interleaved from trial to trial. However, to prevent the proximity effect of the same stimulus presentation, the two presentations of each stimulus were divided into the first and second halves of the task: if a given stimulus was presented with an NA (PA) pair in the first half, it was presented with a PA (NA) pair in the second half.

NA sentences refer to properties of objects presented in the stimulus and thus can be answered from NA. PA sentences concern aspects of conscious experience that can only be captured from PA. In other words, we designed the task to induce NA or PA in the participants using the corresponding pair of sentences. This allowed us to circumvent the notorious difficulty of facilitating that naïve participants, not previously exposed to the phenomenological method, could adopt PA by themselves.

Specifically, the PA sentences consistently inquired about the horizonal structure of the participant’s conscious experience (see Introduction). The fact that conscious experience always involves a horizonal structure was made explicit only after Husserl and other phenomenologists approached conscious experience by adopting the PA. We therefore reasoned that our participants would need to adopt the PA to acknowledge the horizonal aspects of their experience. Accordingly, if they answered correctly to the task in the PA conditions, they would have adopted the PA during the task; further, if they answered correctly in the NA conditions, they would do so while maintaining their NA.

The PA sentences had six categories, each describing a different aspect of the experience horizon: Sentences that describe (i) the ground under a figure; (ii) the pattern that extends beyond an image; (iii) a hidden aspect of an object; (iv) the occluded space between an object and the background; (v) illumination not directly appearing in the image; (vi) perspective onto objects in the image. The NA sentences comprised eight categories, each describing a different aspect of objects in the stimulus or the stimulus itself: Sentences that describe (i) the number of objects of the same color, (ii) the number of objects with the same shape, (iii) the number of objects of the same kind, (iv) the existence of an object with a specific color, (v) the existence of an object with a specific shape, (vi) the existence of an object of a specific kind, (vii) properties of an object or the image, (viii) categories of an object or the image.

We diversified the NA sentences so that participants could engage different cognitive functions, such as counting, visual search, and categorization, while performing the task. This allows us to make sure that the behavioral and neuroimaging data obtained from the NA condition are associated with NA, rather than a specific cognitive function.

### MRI data collection

Functional and anatomical MRI data were collected using a 3T Siemens MAGNETOM Vida scanner (Siemens, Germany) with a 64-channel Siemens volume coil. Functional data were collected during the NPAS task using a multiband gradient echo EPI sequence (26) with the following parameters: repetition time [TR] = 2,000 ms; echo time [TE] = 30 ms; flip angle = 75°; voxel size = 2 × 2 × 2 mm; matrix size = 100 × 100; the number of slices = 75; multiband factor = 3. Anatomical data were collected using a T1-weighted MPRAGE sequence with the following parameters: TR = 2,530 ms; TE = 3.26 ms; flip angle = 9°; voxel size = 1 × 1 × 1 mm; matrix size = 256 × 256; number of slices = 208.

### MRI data preprocessing

MRI data were preprocessed using the standard pipeline of fMRIPrep v20.2.7 (27). T1-weighted images were corrected for bias field inhomogeneities (28) and skull-stripped with ANTs v2.3.3 (29). Brain tissue segmentation of cerebrospinal fluid, and white and gray matter was performed on brain-extracted T1-weighted images using FSL v5.0.9 (30). Brain surfaces were reconstructed and the previous brain segmentation was refined using FreeSurfer v6.0.1 (31). Volume-based spatial normalization to an Montreal Neurological Institute (MNI) standard space was performed through nonlinear registration with ANTs, using the ICBM 152 Nonlinear Asymmetrical template v2009c (32). Functional data were corrected for slice timing (33), head motion (34), and susceptibility distortion (35) in their original, native space. These functional data were then co-registered to anatomical data using FreeSurfer, which implements boundary-based registration (36). Finally, they were resampled into the MNI standard space.

### Searchlight decoding

Prior to whole-brain searchlight decoding analysis (37) with the preprocessed fMRI data in the MNI standard space, several additional preprocessing steps were performed, including linear trend removal, high-pass filtering (128 s per cycle), and z-scoring within each run. Trials in which participants made a wrong choice or provided no response were discarded from the analysis.

This study sought to explore the neural signature of PA toward perceptual experiences by investigating brain activity while participants placed their attention on perceptual experiences under PA compared with NA. In the NPAS task, such brain activity was collected during the 16-s period of the second presentation of an image stimulus in each trial, during which participants observed the stimulus after changing their mode depending on the PA or NA sentences.

To extract brain activity during the second stimulus presentation, the BOLD signals during this period in each trial were transformed into parameter estimates (z-scored β values) by fitting a general linear model (GLM) (38). The model contained nuisance regressors of six head-motion parameters, framewise displacement, and six initial aCompCor components. The reaction times in individual trials were also added to the model as a regressor to regress out the effect of the reaction-time difference between task conditions. This procedure yielded β estimate maps for the individual faces used in the following multivariate pattern analysis.

Then, whole-brain searchlight decoding was employed to explore the cortical localization of stimulus-evoked multivoxel activation patterns distinguishing the NA and PA conditions. While a sphere searchlight with a radius of 6 mm was systematically moved throughout the gray matter, decoding accuracy was quantified on each searchlight location. A linear-kernel support vector machine classifier was trained and tested using a leave-one-run-out cross-validation procedure. In each fold, the classifier was trained using data samples from seven of eight runs (blocks) and then tested on data samples from the remaining run to obtain the decoding accuracy. Training and test procedures were performed independently for each participant to obtain her/his whole-brain map of decoding accuracy.

Next, second-level analysis was performed to examine whether the group-level decoding accuracy was significantly higher/lower than the chance level (i.e., 0.5) on each searchlight location. The significance was evaluated using a one-sample t-test, based on a relatively strict statistical threshold of p < 0.05 after correction for multiple comparisons using FWE at the voxel level. Note that spatial smoothing was employed in neither first-nor second-level analyses.

To examine potential confounding factors in PA-NA decoding, two additional variations of whole-brain searchlight decoding were conducted. In the first, brain activity was decoded to differentiate between trials with slower and faster reaction times. Same as in the original decoding procedure, BOLD signals during the 16-s period of the second stimulus presentation were first transformed into β estimates using a GLM, except that reaction times in individual trials were not included as regressors in this decoding. Subsequently, trials were divided into two groups based on reaction times, and the localization of multivoxel β patterns that distinguished trials with slower and faster reaction times was examined.

In the second variation, brain activity was decoded to distinguish the number of characters in each pair of sentences presented on each trial. In this case, the original method was employed to transform the BOLD signals during the second stimulus period into β estimates through a GLM. Subsequently, after trials were divided into two groups based on the total character count of the pair of sentences presented in each trial, we examined the localization of multivoxel β patterns that differentiated between these two groups of trials. Training and testing procedures for these two variations of decoding analysis remained consistent with those in the original decoding analysis.

### ROI

For further analysis of the neural signals underlying NA and PA, ROIs corresponding to voxel clusters distinguishing between NA and PA conditions were determined according to the second-level decoding analysis. Among voxels with significant t values in the second-level analysis (p < 0.05, voxel-level FWE corrected), separate regions with ≥450 connected voxels (i.e., 3600 mm^3^) were extracted using the function connected_regions in Nilearn (39) and considered as ROIs.

### Analysis of activity strength

The decoding analysis allowed us to identify brain regions that contain information distinguishing NA and PA but not to understand changes in the activity strength of the regions depending on the attitudes. Therefore, we used the extracted ROIs for examining changes in the activity strength of the ROIs. In each condition, parameter estimates (z-scored β values) obtained from the GLM prior to decoding analysis were averaged over all voxels within each ROI to yield the estimated activity strength in that ROI for each participant. Then, the estimated activity strength of each ROI in the NA condition and that in the PA condition was compared across participants using a paired t-test with ROI-level FWE correction for multiple comparisons.

## Acknowledgments

We thank Ms. Hitomi Koyama for her experimental support. This work was supported by JSPS KAKENHI Grant-in-Aid for Scientific Research B (21H03535 to SN), Scientific Research C (21K00011 to TN, SN, HTH, and KM and 20K00001 to KM, SN, HTH, and TN), Exploratory Research (22K19819 to SN), and Transformative Research Areas B (24H00810 to SN, 24H00807 to SN and TK, and 24H00808 to TK).

## Footnotes

1 Recent studies show that affordance perception can take place with or without motivation to realize the afforded action [e.g., (16, 17)]. Accordingly, some think that our sensitivity to affordances consists of two layers, (i) perception of an affordance and (ii) potentiation (or automatic preparation) of the afforded action (18). Accordingly, we might explain the inhibited activity in the cerebellum during PA as a result of the inhibition of potentiation of the afforded action during PA. Generally, this distinction could allow us to develop a more fine-grained explanation of brain activities corresponding to PA identified in our neuroimaging data.

## Supporting Information

**Fig. S1.**
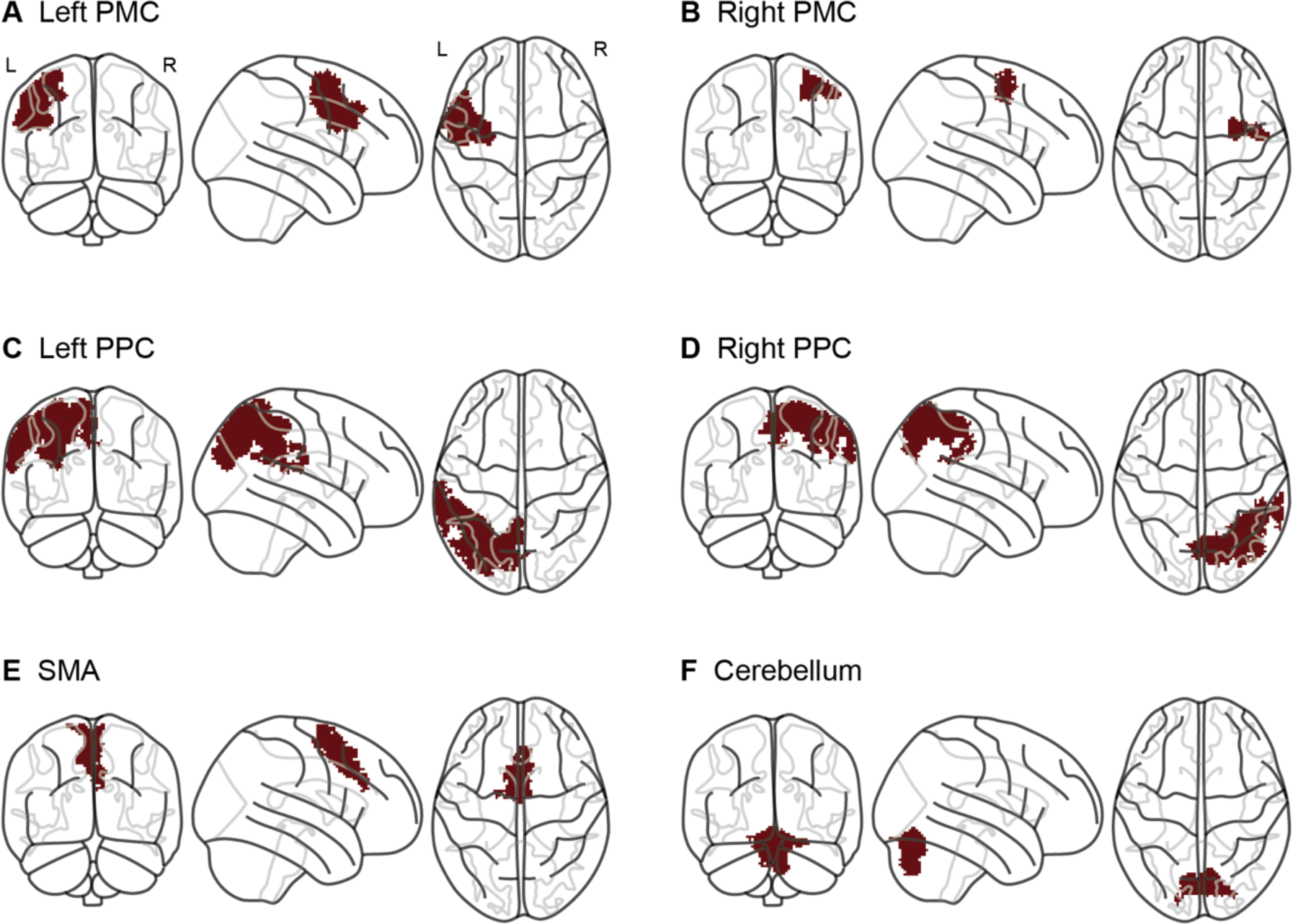
Six brain regions used for ROI analysis. Six brain regions corresponding to voxel clusters distinguishing NA and PA conditions were defined as ROIs according to second-level decoding analysis. Each ROI included connecting significant voxels forming a volume above a predefined threshold (3600 mm^3^; for more details see Materials and Methods). The six ROIs, indicated by brown-filled areas, were the bilateral PMC (A and B), the bilateral PPC (C and D), the SMA (E), and the cerebellum (F). See also SI Appendix, Table S1.

**Fig. S2.**
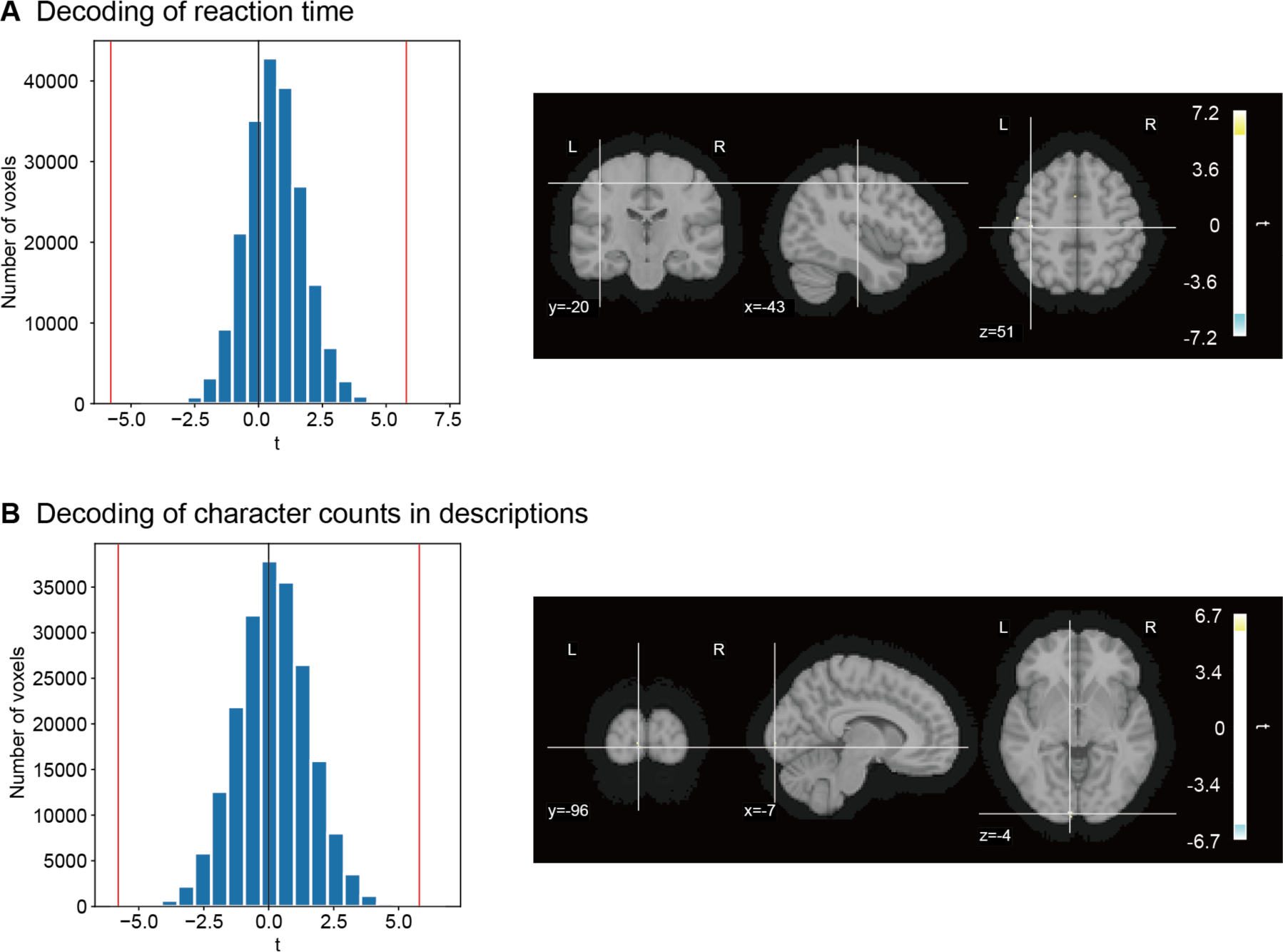
Whole-brain searchlight decoding to identify brain regions with changing multivoxel activation patterns due to confounding factors. (A) Decoding of reaction time. Two groups of trials equally divided according to reaction time were classified from multivoxel activation patterns (for more details see Materials and Methods). The decoding accuracy, transformed into t statistics, is presented in a histogram (left) and in brain slices (right). In the histogram, vertical red lines indicate the significant threshold; the same as used for main decoding analysis (Fig. 3; P < 0.05, voxel-level FWE corrected). In brain slices, colored locations represent the centers of searchlights exhibiting significant decoding accuracy. Only six voxels showed significant decoding accuracy. (B) Decoding of character counts in sentences. The two groups of trials equally divided according to the total character counts in two sentences on each trial were classified from multivoxel activation patterns (for more details see Materials and Methods). The conventions are the same as in A. Only eight voxels showed significant decoding accuracy.

**Fig. S3.**
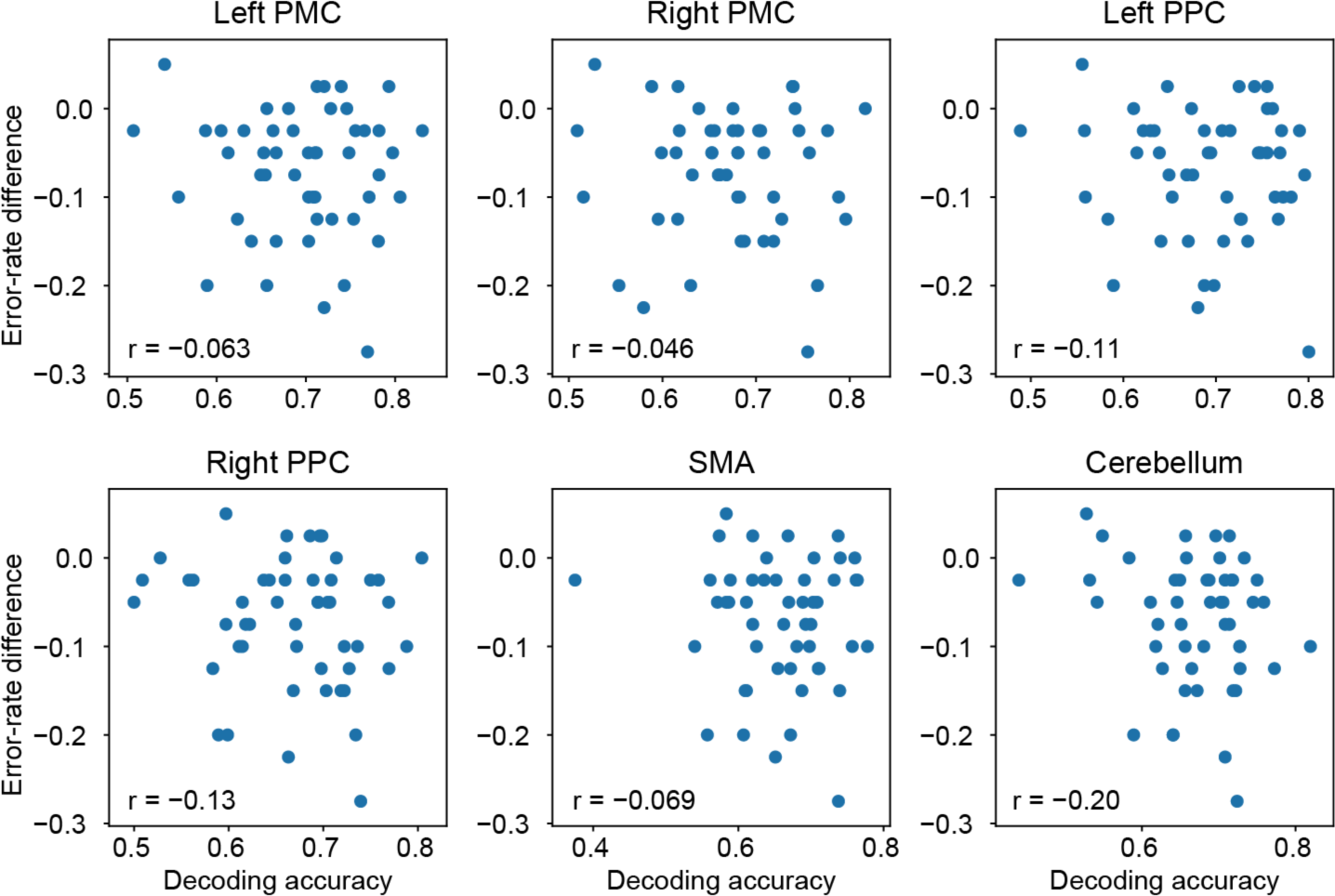
No relationship between decoding accuracy and between-condition differences in error rate. The difference in error rates between PA and NA conditions (Fig. 2) raises the possibility that this error-rate difference might cause changes in brain responses between conditions. To investigate this, we examined whether there is a correlation between error-rate differences and decoding accuracy among participants in each brain region used for ROI analysis (SI Appendix, Fig. S1). Each panel shows the decoding accuracy (x-axis) and error-rate differences (y-axis) for individual participants in each region; each dot depicts a single participant. We found no significant correlation any region (P > 0.17, uncorrected), indicating that the observed changes in brain responses are unlikely to be explained by error-rate differences.

**Table S1.** Details of brain regions used for ROI analysis.

|  | MNI coordinates |  |  | Volume<br>(voxels) | Peak t value |
| --- | --- | --- | --- | --- | --- |
|  | x | y | z |  |  |
| Left PMC | −46 | 4 | 42 | 2282 | 13.28 |
| Right PMC | 36 | −1 | 56 | 483 | 9.77 |
| Left PPC | −27 | −65 | 51 | 4809 | 10.40 |
| Right PPC | 27 | −62 | 56 | 2939 | 10.77 |
| SMA | 0 | 4 | 60 | 1136 | 11.16 |
| Cerebellum | −6 | −64 | −35 | 6631 | 5.77 |

MNI coordinates indicate the voxel with the peak t value for each ROI. See also SI Appendix, Fig. S1.

**Dataset S1.** All image stimuli and the corresponding PA and NA sentence pairs used in the NPAS task. ***This file is not included in this preprint version of the manuscript to avoid the inclusion of photographs and any other identifying information about people*.**

